# Structural Basis of Mutation-Dependent p53 Tetramerization Deficiency

**DOI:** 10.1101/2022.05.13.491836

**Authors:** Marta Rigoli, Giovanni Spagnolli, Giulia Lorengo, Paola Monti, Raffaello Potestio, Emiliano Biasini, Alberto Inga

**Author notes:** Authors contributed equally to this work (Marta Rigoli, Giovanni Spagnolli). Correspondence could be addressed to, or. Sibylla Biotech SRL, Piazzetta Chiavica 2, 37121 Verona.

## Abstract

The formation of a tetrameric assembly is essential for the ability of the tumor suppressor protein p53 to act as a transcription factor. Such a quaternary conformation is driven by a specific tetramerization domain, separated from the central DNA binding domain by a flexible linker. Despite the distance, functional crosstalk between the two domains has been reported. This phenomenon can explain the pathogenicity of some inherited or somatically acquired mutations in the tetramerization domain, including the widespread R337H missense mutation occurring in the population of south Brazil. In this work, we have combined computational predictions through extended all-atom molecular dynamics simulations with functional assays in a genetically defined yeast-based model system to reveal structural features of p53 tetramerization domains and their transactivation capacity and specificity. Besides the germline and cancer-associated R337H and R337C, other rationally designed missense mutations targeting a significant salt bridge interaction that stabilizes the p53 tetramerization domain were studied (R337D, D352R, and the double mutation R337D plus D352R). Simulations revealed a destabilizing effect of pathogenic mutations within the p53 tetramerization domain and highlighted the importance of electrostatic interactions between residues 337 and 352. The transactivation assay performed in yeast by tuning the expression of wild-type and mutant p53 proteins revealed that p53 tetramerization mutations could decrease transactivation potential and alter transactivation specificity, in particular, by better tolerating the negative features in weak DNA binding sites. These results establish the effect of naturally occurring variations at positions 337 and 352 on p53 conformational stability and function.

## INTRODUCTION

The *TP53* gene encoding the well-known tumor suppressor protein p53 is undoubtedly one of the most critical cancer genes ^1^. Acting primarily as a nuclear sequence-specific transcription factor, P53 coordinates a complex network of gene targets to modulate many cellular pathways that respond to many cellular stress conditions ^2^. The p53 transcriptional network is pleiotropic. The responses are distributed to many distinct target genes that collectively determine a specific cell outcome among the many possible, including cell cycle arrest and cell death as archetypes ^3^. Furthermore, the p53-controlled pathway is highly regulated by multiple feedback loops at transcriptional, post-transcriptional, and translational levels ^4,5^. This complexity is exemplified by the notion that there is still a significant lack of knowledge about the critical tumor-suppressive functions of p53 in specific cancer types or evolutionary cancer tracks ^6–8^.

The p53 function as a transcription factor relies on the assembly of a p53 tetramer that can interact with variations of DNA target sites known and referred to as Response Elements (REs) ^9–11^. The p53 tetramer conformation results in a dimer of dimers, with p53 dimers being formed co-translationally ^12^. This assembly is made possible by the presence and features of a so-called oligomerization or tetramerization (TET) domain in the proximal carboxy-terminal region (C-ter) of the protein ^11,13^. The structure of the human p53 TET domain has been resolved by crystallography and NMR and consists of a four-helix bundle, with each p53 monomer featuring a sheet-loop-helix fold ^14,15^. p53 tetramer assembly and stability are also influenced by interactions between the DNA binding domains (DBD). These are thought to be modulated by long-range interactions between the distal carboxy-terminal region of the protein and the DBD ^15–17^. The transactivation domains are critical for p53 function ^18,19^ and are located in the amino-terminal sequence (within the first 60 amino acids), a region that is considered to be highly unstructured ^20^. Together with the last 30 amino acids in the C-ter, it is subject to several post-translational modifications that impart changes in protein-protein interactions, conformation, subcellular localization, and function to the protein ^21,22^.

The p53 binding site in the DNA reflects and co-evolved with the quaternary conformation of the protein. As the active protein structure is a p53 tetramer, so the p53 REs comprises an arrangement of four binding sites, consisting of 5 nucleotides, which are organized as inverted repeat dimers (i.e., a half-site), and two such dimers are placed adjacent or closely spaced to form an entire site ^23–28^. Many studies based both on *in vivo* approaches, such as ChIP-seq or ChIP-Exo, and *in vitro* assays, such as Selex-seq, have derived the features of the p53 consensus RE and cataloged the many variants present in genomes (reviewed in ^26,28,29^). Further, biochemical, biophysical, or defined gene reporter assays have deconstructed the p53 RE to reveal its critical features ^23,25,29^. These studies also established the consequences of polymorphisms or mutations in the p53 REs or the p53 protein itself, which are among the most frequent somatic alterations acquired by human cancer ^30–34^. These studies also established that p53 tetramers could be assembled on DNA as dimers, and p53 tetramers can also bind hemi-specifically, with one dimer establishing sequence-specific contacts with a decamer binding site and the second p53 dimers contacting non-specifically the DNA backbone ^13^.

Interestingly, there is clear evidence not only that the p53 RE is highly degenerated, which results in many different versions of the binding site enabling various levels of quantitative controls by p53, but also there has been an apparent selection pressure during evolution to avoid optimal p53 binding sites in promoters that would lead to constitutive, non-regulatable p53-dependent transcriptional control of a target gene^23,34,35^.

Given the essential requirement for the tetrameric structure, it is not surprising that the TET domain has undergone a significant evolutionary divergence within the p53 family of proteins that also comprises p63 and p73 ^36–39^. Also, within p53 protein from different and distant species, there is evidence of a divergence in the features of the TET domain that can also impact how the p53 tetramer interacts with the DNA, particularly for what concerns the relative arrangements of the two DNA half-sites ^40,41^.

Within the spectrum of p53 mutations observed in human cancers, those at the TET domain are much less abundant than mutations in the central DBD of the protein; in fact, nearly 80% occur at residues in this latter domain ^42^. p53 mutations are among the most frequent somatic alterations occurring in human cancer ^1^; furthermore, p53 mutations can also be inherited in the germline, where heterozygous mutant alleles are associated with a highly penetrant cancer predisposition syndrome ^43,44^. Also, in the case of germline p53 alleles, there is a strong enrichment for missense mutants in the p53 DBD ^45^.

The low frequency of mutations in the p53 TET domain can be explained, at least in part, by the fact that p53 missense mutations in the DBD can also acquire oncogenic gain of function (GoF) ^46,47^. GoF, however, requires that the mutant protein remains able to be localized in the nucleus and participate in protein complexes, modulating aspects of DNA replication or transcription. Conversely, an intact P53 tetrameric conformation is critical to masking a nuclear export signal. However, tetramerization can enable the occurrence of mixed tetramers comprising WT and mutant DBDs in cells undergoing a transformation that has acquired a heterozygous p53 mutation, a feature of mutated alleles frequently referred to as dominant-negative ^48,49^. Finally, systematic mutagenesis of the human p53 TET domain based on single nucleotide changes showed that most of those mutants would not dramatically reduce p53 transactivation function in a reporter assay ^50^.

There are, however, a few exceptions to the general rule of the scarcity of p53 TET domain mutations in cancer ^51^. Two missense changes in the p53 TET domain have been found in the germline (R337H and R337C, numbering of the human protein), affecting a conserved arginine involved in establishing a salt bridge with D352 that is important for the stability of the dimeric conformation (Figure 1). While germline R337C is associated with a classical tumor proneness spectrum, R337H was initially identified as predisposing to pediatric adrenocortical carcinoma ^52,53^. Molecular epidemiology studies, a screening campaign, and a knock-in mouse model have clarified that this allele results in a partially functional protein whose degree of inactivation can be related to pH and the protonation level of the histidine residue ^54–56^. The allele is widespread in the population in southern Brazil. It is associated with variable penetrance to various pediatric and adult-onset cancers, including choroid plexus carcinomas and breast cancer ^53^.

**Figure 1.**
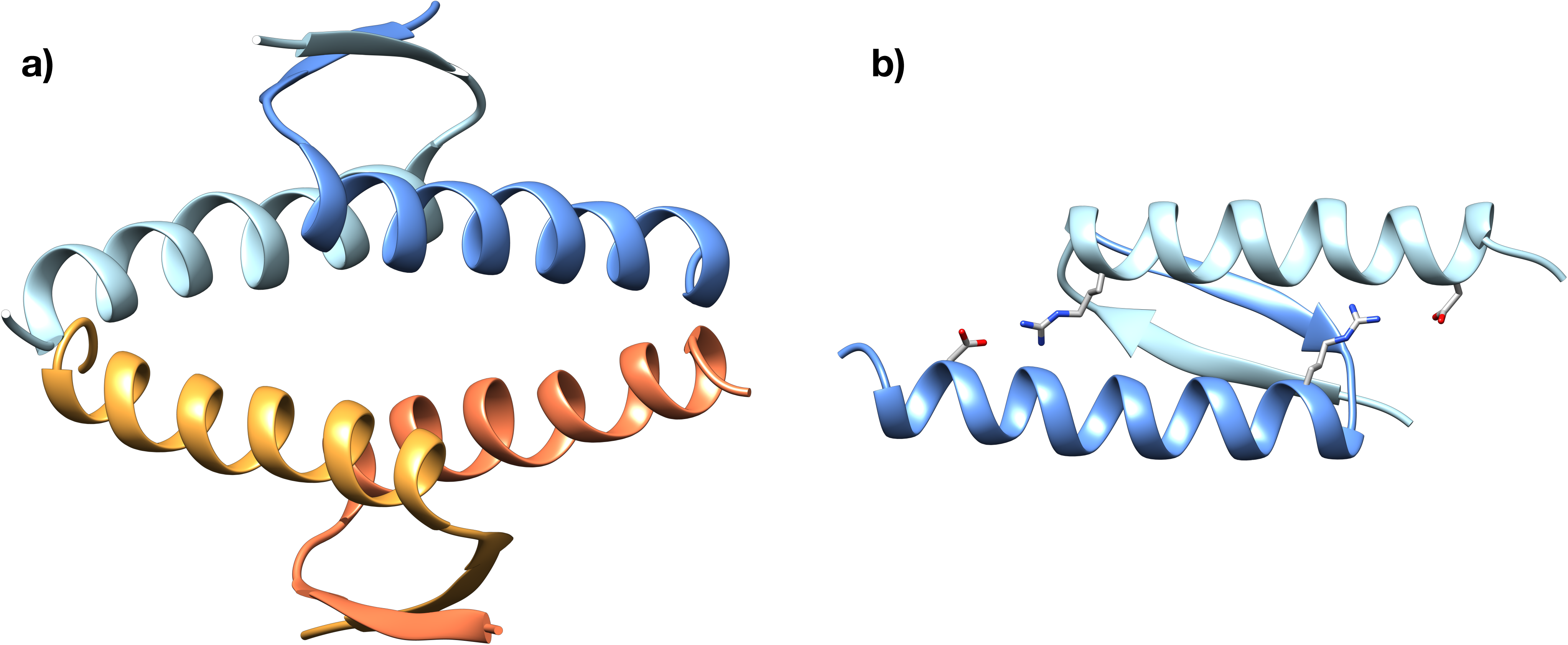
Structure of the p53 TET domain. **A**. Ribbon representation of the fully assembled TET domain of WT p53. Individual chains are depicted in different colors. **B**. Representation of a dimer of the p53 TET domain displaying the two intermolecular salt bridges between residues R337 and D352.

Given the centrality of p53 in cell biology, the strong selection to lose or at least attenuate p53 responses in cancer, and the fact that mutant p53 proteins are over-expressed in cancer cells, the potential to reactivate p53 mutant proteins by pharmacological strategies as cancer therapy has a robust rationale and was pursued in many ways ^57–59^. Cell-based assays have identified potential small molecules of therapeutic interest, but it has proven difficult to establish the exact mechanism of action of the hits coming from these assays ^60^. Potent rationally designed molecules have also been developed primarily targeting the interaction between p53 and its major negative regulator MDM2 but have shown high toxicity in clinical trials ^61^. Molecules patching the thermodynamic instability caused by specific p53 mutations, such as Y220C, or trying to increase the stability of the p53 DBD have also provided proof of concept evidence of the value of rescuing p53 folding ^62^. The lack of a high-resolution structure of the full-length p53 tetramer bound to DNA represents a limitation in rational drug design. With a few exceptions, all the effort has been directed to p53 mutant proteins in the DBD ^63–65^. Molecules targeting the TET domain could be highly valuable, especially for individuals who have inherited the R337H mutations ^66^, but in principle also for other classes of cancer-associated p53 alleles in the DBD that retain partial function and whose defect could, in principle, be compensated by increasing the stability of the tetramer conformation.

In silico modeling, particularly molecular dynamics (MD), has been attempted to boost the search for small molecule leads to rescue p53 mutations ^67^. Still, until recently, those approaches have been somewhat limited by the relatively short simulation times that could be afforded.

Here we have adopted a state-of-the-art Molecular Dynamics approach to study p53 TET mutant proteins R337C and R337H along with rationally designed mutations to probe the impact of the salt bridge between R337 and D352 on the stability of the p53 TET domain and the consequences not only on transactivation potential but also on transactivation specificity. Indeed, by coupling computational predictions through extended all-atom molecular dynamics simulations with the results of a genetically defined transactivation-based assay in yeast, we reveal that targeting the R377-D352 salt bridge interaction in the p53 TET domain can lead to reactivation of p53, but also an apparent change in relative transactivation specificity.

## RESULTS

### 1. Computational prediction of the effect of naturally occurring or rationally designed *TP53* mutation on the stability of the p53 TET domain

Mutations within the TET domain of p53 have been associated with Li-Fraumeni or Li-Fraumeni-like cancer predisposition syndromes. These mutations are thought to affect the stability of the tetramer by altering the intermolecular salt-bridge formed by residues R337 and D352 (Figure 1). We sought to test this possibility by performing extended, all-atom molecular dynamics simulations. First, we simulated for 2 µs the wild-type (WT) p53 TET domain and the two disease-associated mutant proteins (R337C and R337H). We included three different states of His337, accounting for the protonation of δ (R337H_δ_), ε (R337H_ε_), or both (R337H_δε_) imidazole nitrogens. Next, we simulated 2 µs of three artificial p53 mutant proteins, designed to test the contribution of the intermolecular electrostatic interaction between residues 337 and 352 (D352R, R337D, and R337D/D352R). Our simulations indicated that disease-associated substitutions at residues 337 (R337H_δ_, R337H_ε,_ and R337C) destabilize the conformation of the p53 TET domain. Interestingly, the R337H_δε_ form, which carries a positive charge on the side chain similarly to its WT counterpart, showed no stability alterations (Figure 2A). Artificial p53 mutant proteins showed differential destabilization patterns, with the R337D exhibiting the most prominent alteration, the R337D/D352R a more modest effect, while the D352R resembled the WT form (Figure 2B).

**Figure 2.**
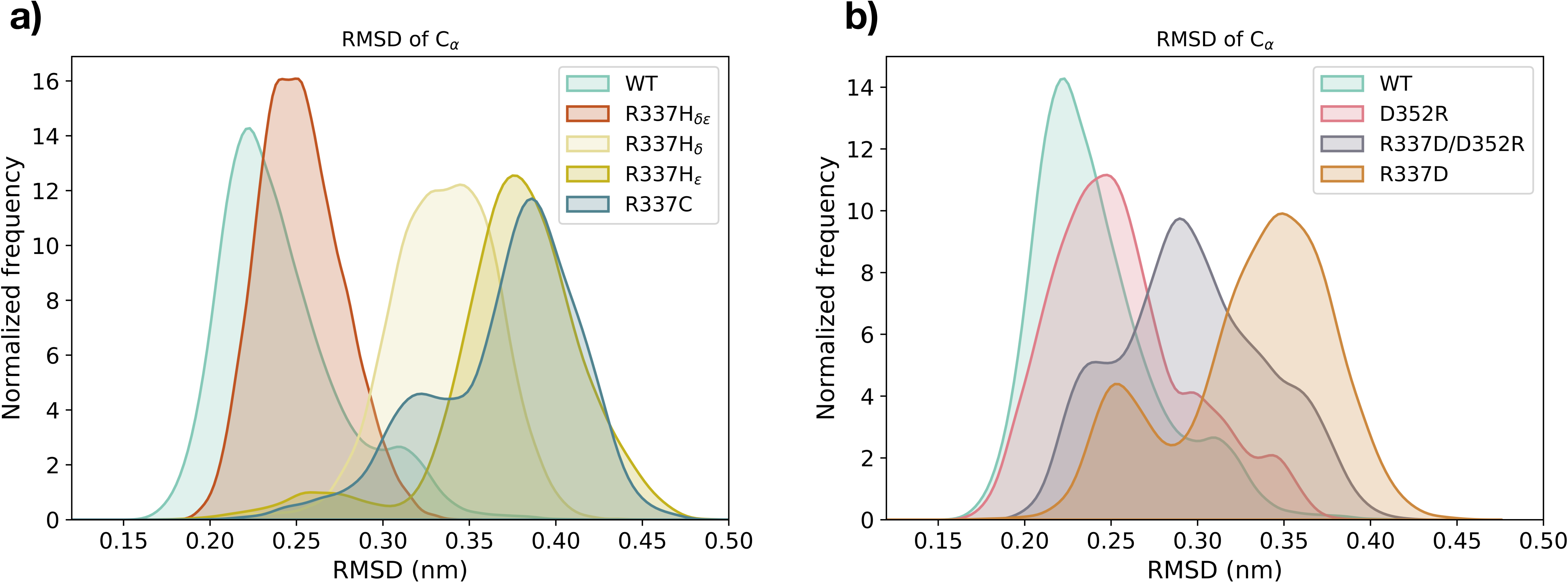
Molecular dynamics simulations of p53 TET domain mutants. **A**. The destabilization effect of disease-associated mutations lying within the p53 TET domain was analyzed by molecular dynamics. The graph shows the RMSD distribution of the WT tetramer and the R337C and R337H mutants. The latter was simulated in three different states of the histidine, including the protonation of δ (R337H_δ_), ε (R337H_ε_), or both (R337H_δε_) nitrogens. The results indicate that the WT and R337H_δε_ show overlapping distributions with low RMSD values. Conversely, the R337H_δ_, R337H_ε_, and R337C mutants exhibit distributions centered at higher RMSD values. **B**. The effect of rationally-designed substitutions, including D352R, R337D, and R337D/D352R, on the stability of the p53 TET domain was compared to the WT counterpart. The results indicate that the WT and the D352R form show distributions with low RMSD values. Conversely, the R337D/D352R and R337D exhibit distributions centered at higher RMSD values, with the latter presenting a more prominent destabilization. Each system was simulated for 2 µs.

To corroborate these data, we computed the inter-chain distances of residues 337 and 352 within the p53 tetramer for all the simulated systems. The results show that, similarly to the WT, a charged histidine at position 337 and the double mutant protein R337D/D352R display distance distributions compatible with the formation of stable salt bridges. Close interaction distances are also observed for R377-D352 in the D352R mutant, reflecting π-π and π-cation interactions between the guanidinium groups of the arginines. Conversely, in all the other conditions, a distance incompatible with a stable interaction was detected in at least two pairs of residues (Figure 3).

**Figure 3.**
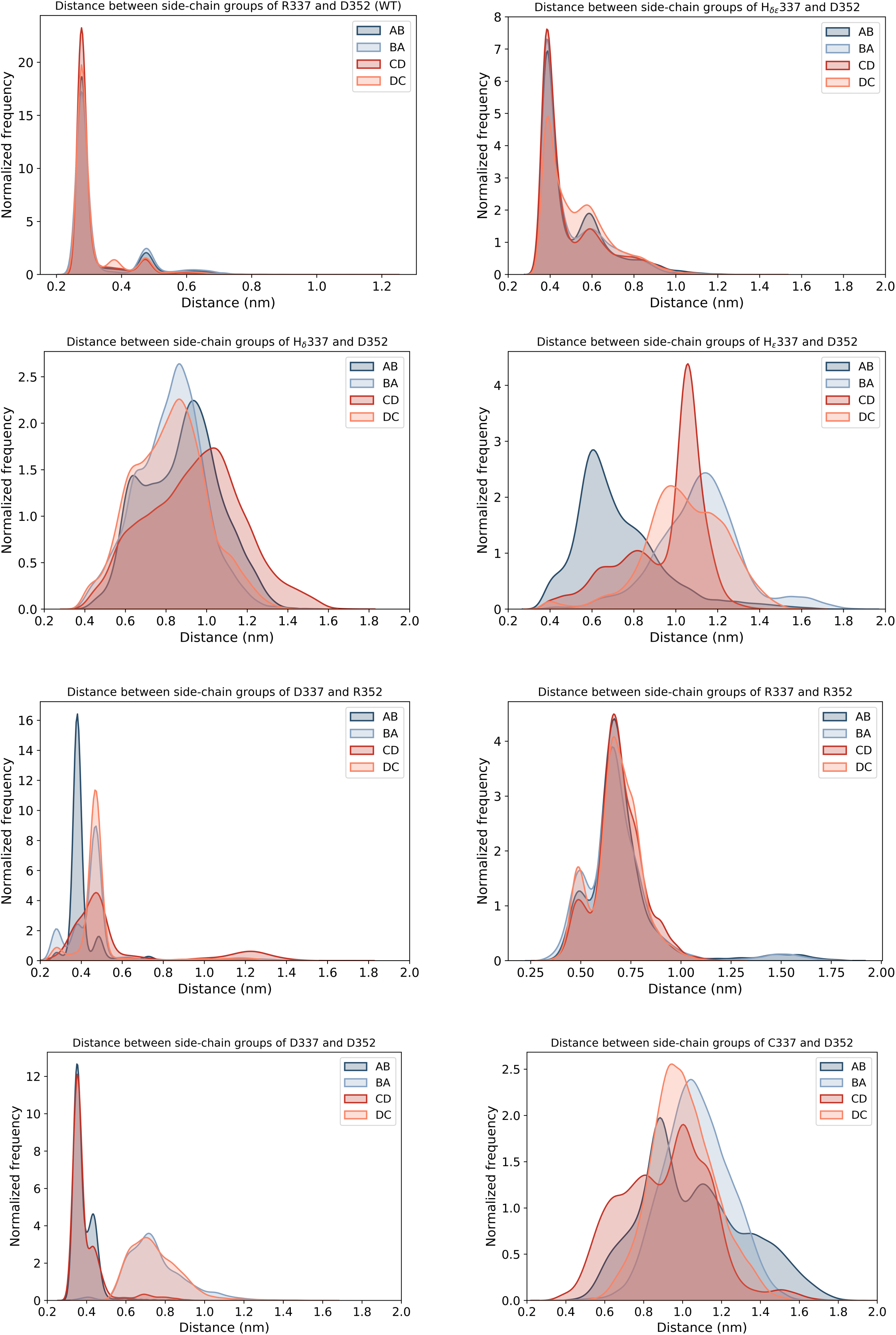
Analysis of the distances between side chains of residues 337 and 352 within the p53 tetramer. The inter-chain distance between residues involved in the salt bridge formation at positions 337 and 352 in the WT protein was analyzed for all the simulated systems. The graphs display the measure of the distance between the centers of mass of the functional groups of each couple of amino acids (337 and 352), initially forming salt bridges between two pairs of the four polypeptide chains of the tetramer (A with B and C with D), resulting in the 4 different arrangements (337.A-352.B, 337.B-352.A, 337.C-352.D, and 337.D-352.C). The results show that only the charged histidine at position 337, the double mutant R337D/D352R, and the D352R mutant display distance distributions compatible with the formation of effective interactions.

The alteration of an intermolecular interaction resulting from an amino acid substitution may destabilize regions in the protein complex lying several residues apart. To assess this possibility for residues 337 and 352 of the p53 TET domain, we computed the average contact map differences between the trajectories of the WT and the mutant forms. As expected, the contact maps of D352R and R337H_δε_ displayed no substantial deviation from the WT. Conversely, R337C, R337D/D352R, R337D, R337H_ε,_ and R337H_δ_ showed a significant variation, with the latter presenting the most prominent effects (Figure 4).

**Figure 4.**
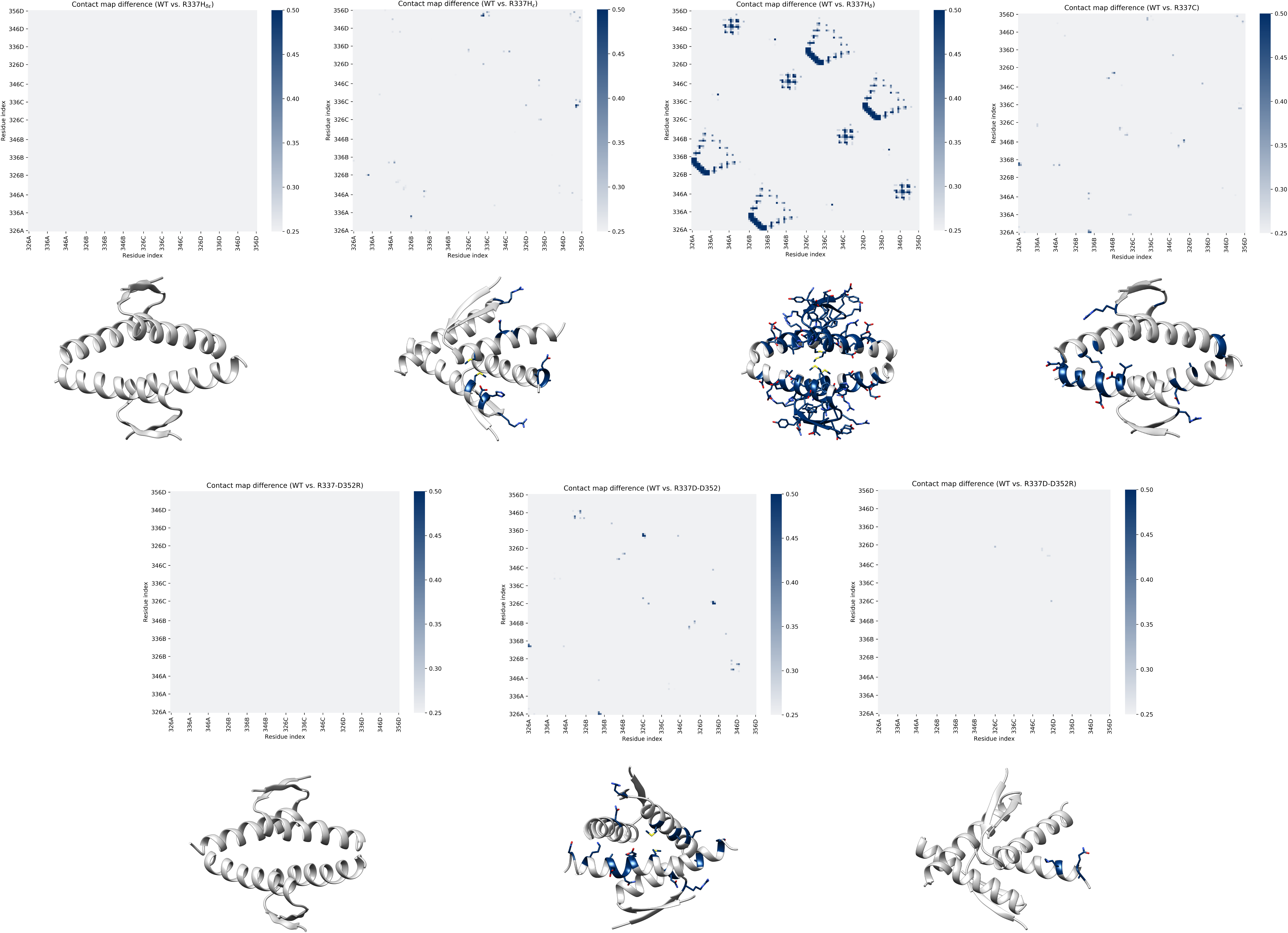
Contact map analysis of WT and mutant p53 TET domain. The average contact map difference is represented as a residue index matrix for each form. Mutant p53 forms are expressed as the absolute value of the difference with the WT protein. The contact maps of R337C, R337D/D352R, R337D, R337H_ε_, and R337H_δ_ show a significant variation, with the latter presenting the largest divergence. In contrast, D352R and R337H_δε_ display no substantial deviation from the WT. The side chains of residues with the most evident differences in each contact map, defined with a threshold between 0.25 and 0.5, are illustrated as sticks.

In summary, these results define the mechanism by which residue variations in positions 337 and 352 destabilize the TET domain of p53 and predict a structure-function relationship.

### 2. Experimental analysis of mutation-dependent, p53-driven transcription

Based on the results of the MD simulations, we designed a set of experiments to validate the predicted effects of each p53 mutant protein. The goal was achieved by taking advantage of a defined functional assay in yeast where p53 alleles can be expressed at variable levels under an inducible, finely tunable promoter, and the p53 transactivation potential is quantified based on the level of activation of the firefly reporter gene that is cloned in a single copy at a specific chromosomal location. A panel of reporter strains that are entirely isogenic except for the sequence of the p53 RE driving p53-dependent reporter gene transactivation are used to establish the consequences of p53 alleles on sequence-specific transactivation specificity. Specifically, we tested WT p53, the naturally occurring R337C and R337H alleles, and the ad-hoc designed mutations R337D, D352R, and the double mutant R337D/D352R. The relative transactivation of each protein was determined by culturing yeast reporter strains stably transformed with centromeric p53 reporter plasmids and cultured in media containing different amounts of galactose to vary the level of expression of p53 proteins. Results confirmed that R337C is a near loss-of-function mutation. At the same time, R337H exhibits a more subtle transactivation defect that is appreciable, particularly when p53 protein levels are low (Figure 5, Figure S3), consistent with previous studies. R337D is a complete loss of function allele, while D352R is WT-like. The double mutant R337D/D352R shows only a partial rescue of the R337D effect (Figure 5, Figure S3). Interestingly, we noticed that the relative activity of this panel of mutations was affected by the level of expression of the p53 alleles and by the nature of the p53 RE. We tested two REs derived from the very well-established p21 target gene; one RE is the natural sequence derived from the human promoter and known as p21-5’, while the second is derived from p21-5’ but contains a two-nucleotide spacer between the decameric half-sites (p21-SP2). Consistent with previous studies, we confirmed that the small spacer significantly diminished p53 transactivation potential and required higher galactose levels to measure p53-dependent transactivation (Figure 5 and Figure S3). Unexpectedly, however, the presence of the spacer also led to a change in the relative activity of the p53 TET domain. In particular, D352R showed much higher transactivation potential than WT p53, and, consistent with that observation, the double mutant R337D/D352R was fully rescued. Also, the R337H allele was WT-like in the p21-SP2 reporter strain or slightly more active. These differences in relative activity were not solely related to the lower DNA binding affinity of WT p53 for the REs. We also tested the panel of TET mutations with one additional yeast reporter strain based on the p53 RE found in the human PUMA/BBC3 gene. This RE has no spacer between the decameric half-sites, like the canonical p21-5’ RE, but, due to deviations from the RE consensus sequence, it mediates moderate responsiveness to p53. R337C was confirmed as a near loss-of-function allele in the PUMA reporter strain, R337D as a complete loss of function, R337H, and D352R as partial function alleles. The double mutant R337D/D352R was only slightly active (Figure S3). Finally, when p53 expression levels were high in 0.032% galactose (or 0.128% for the p21-SP2 reporter), R337H and D352R were WT-like, and the double mutant was fully or nearly fully rescued. Still, only the structurally altered p21-SP2 reporter showed enhanced activity of D352R. All the above-described differences in transactivation potential were undoubtedly not related to variations in the level of protein expression, as confirmed by western blot (Figure S4).

**Figure 5.**
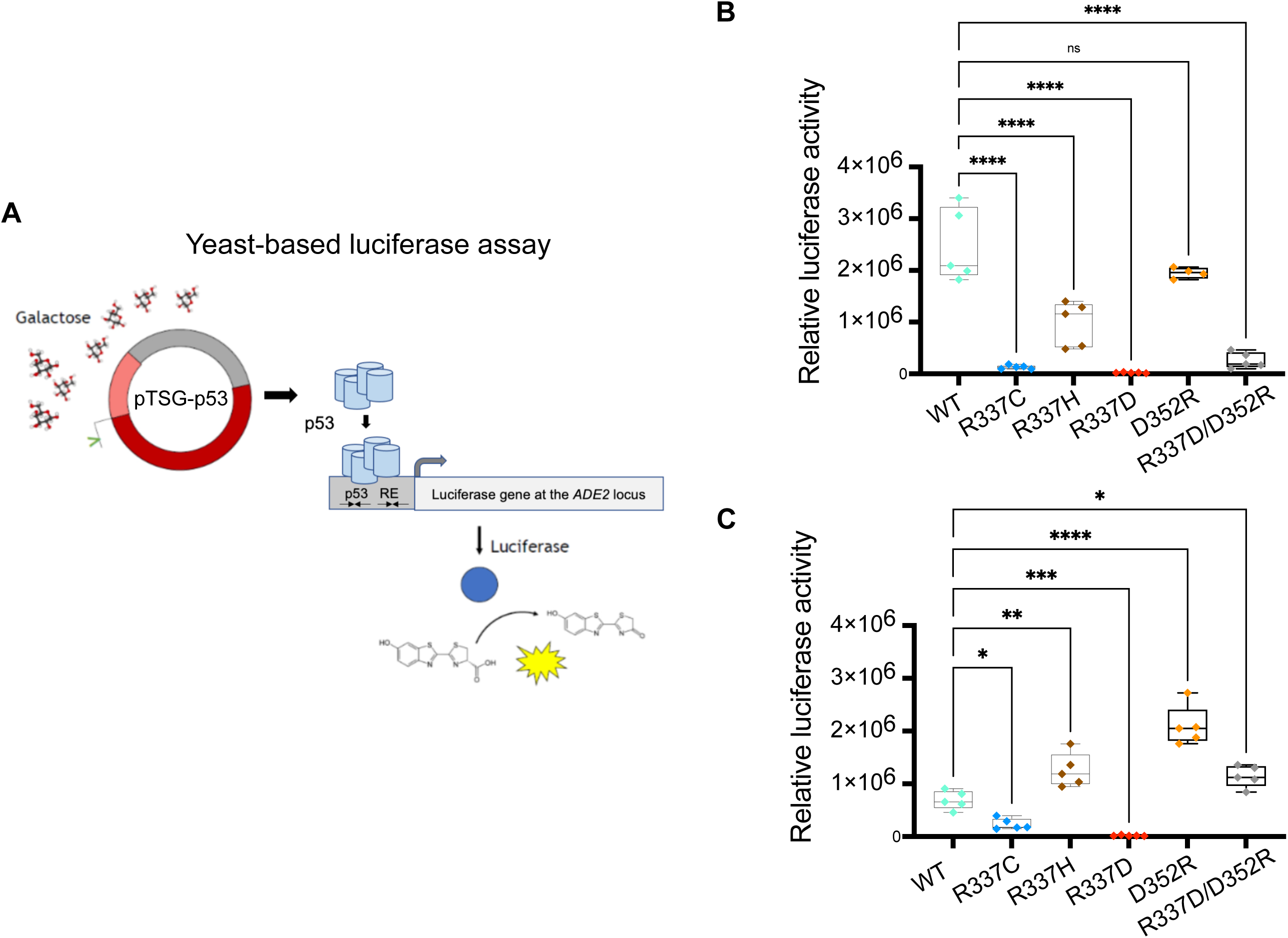
Transactivation potential of p53 TET mutations. **A**. Scheme of the experimental approach. p53 expression is achieved by stable transformation of yeast reporter cells with plasmids that, by containing an origin of replication and a centromeric sequence, ensure stable low-copy transmission to daughter cells. The vector is based on the pRS314 shuttle vector and contains a selectable TRP1 marker gene complementing an auxotrophy of the yeast strain. The vector contains a GAL1,10 promoter driving the expression of the human p53 cDNA. This promoter affords inducible expression that can be modulated precisely by varying the amount of galactose in the medium. The yeast reporter strain contains the *Photinus Pyralis* luciferase cDNA cloned on chromosome XV at the endogenous ADE2 locus. The luciferase transcription is regulated by a minimal promoter that can be stimulated by p53 binding to upstream RE. Permutation of the p53 RE in isogenic yeast strains was made possible by an oligonucleotide targeting approach (see methods for details). A. Top panel; relative transactivation of the indicated p53 alleles expressed at moderate levels by medium containing 0.008% galactose for six hours in a yeast reporter strain containing the high affinity p53 RE derived from the p21 promoter (p21-5’). Luminescence is normalized to the optical density of the cultures. Plots present the average normalized light units, the confidence intervals, and the individual data points after subtracting the p53-independent activity of the reporter. B, Lower panel; same as the top panel except that the experiment was performed using a reporter strain that relies on a p53 binding site derived from the p21-5’ RE but modified by the inclusion of a 2nt spacer between the two decameric half-site motifs. Given the lower responsiveness of the spaced RE, a 0.032% galactose concentration was used. * p<0.05; *** p<0.001; **** p<0.0001, ns = not significant; one-way ANOVA with Dunnett’s multiple comparison test.

## DISCUSSION

High-resolution computational modeling that assesses stability and dynamic folding changes in the p53 protein will be essential not only to reveal the molecular basis of functional alterations caused by disease-associated mutations but, in principle, also to identify structural intermediates that could be qualified as target sites for the design of pharmacological rescue molecules. In the case of p53, the possibility of developing drugs that can restore its conformation or improve its thermodynamic stability would have a high potential to benefit cancer patients ^57–59,61,68,69^. Drugging p53 has, however, proven very difficult not only because the protein is a transcription factor and hence lacks a well-defined catalytic site but also because of the incomplete availability of data on the structure and dynamics of the entire functional unit of the protein, *i*.*e*., a p53 tetramer that is competent to bind to DNA and stimulate transcription. There are crystal structures available for the p53 DNA binding domain, both the WT, alone or attached to different DNA target sites, some hotspot p53 mutations ^11,70–77^, and the p53 TET domain ^14^.

However, there is still a lack of data to fully reveal the structural details of the crosstalk between the different domains, including the highly unstructured and heavily post-translationally modified N-terminal and C-terminal domains ^11,17,78,79^. This lack of knowledge represents a severe limitation as p53 is a highly versatile transcription factor that interacts through a wide range of affinity with many DNA RE binding sites that can be structurally diverse in their internal organization ^13,27^. Hence, while increasing the thermodynamic stability of the p53 DNA binding domain can increase p53 transactivation and rescue the consequences of some cancer-associated mutations ^80–83^, it can also affect transactivation specificity towards the various categories of p53 binding sites ^3,5,28^, with functional effects that are difficult to predict^84^.

Conversely, cell-based assays can select for small molecules that result in a specific p53-dependent cellular outcome ^59,61,85^. Still, given the extreme pleiotropy of p53 and the pluralism of highly connected molecular pathways, it has proven challenging to reveal the precise mechanisms of action on p53 of leads emerging from those screening campaigns. All-atom MD simulations hold great potential to reveal the molecular defect of p53 mutations in terms of local structural distortions, long-range spatial effect, and kinetic consequences on conformational changes ^67,86–88^. MD has been applied to study a panel of hotspot p53 missense mutations in the DNA binding domain ^89^ to model the p53 TET domain. There have been attempts to combine high with low resolution modeling to address the entire p53 tetramer ^86,90^. For example, those studies have suggested the existence of specific structural features that are more evident or stable in the mutant p53 DBD and could be druggable ^89^. In the case of the p53 TET domain, structure and molecular dynamics data have led to the attempt to design molecules that could lead to enhanced stability ^66^. Our results directly suggest that the TET domain and DNA binding domain are not independent, as an alteration in the TET domain could also impact the arrangements of p53 monomers and dimers, which in turn results in a variety of functional consequences. Such information could allow future therapeutic strategies to restore mutant p53 stability in human cancer.

## METHODS

### Molecular dynamics simulations

Starting from the WT PDB structure (PDB code: 2J0Z), the mutant models were built by using the software Chimera ^91^. The proteins were solvated in water using the TIP3P model ^92^; Na and Cl ions were added to neutralize the protein’s charge and mimic the physiological salt concentration (150 mM). The simulation box was chosen for the cubic shape, and the protein had a minimum distance of 1 nm to the box edge. The simulations were performed in the NPT ensemble at 300 K and 1 bar, using the stochastic velocity-rescale thermostat ^93^ and the Parrinello-Rahman barostat ^94^. The chosen parameters are: temperature coupling constant tau-t = 0.1ps, pressure coupling constant tau-p = 2ps. The integration step was set to 2 fs, the selected integrator was the leap-frog one, and the LINCS algorithm ^95^ was selected to apply holonomic constraints. The forcefield employed for the protein simulations is the Amber14sb ^96^.

### Yeast cultures

The yeast reporter strains were previously developed from the yLFM-CORE strain and the so-called Delitto Perfetto targeting strategy ^97^. Cells were transformed with centromeric plasmids containing the expression cassette for human p53 alleles ^30^. The promoter is derived from the *GAL1*,*10* gene, and we established it could be regulated by varying the sugar concentrations in the culture medium. Specifically, the expression of p53 is repressed in glucose media, reaches a slightly higher basal level when glucose is replaced by galactose, and can then be gradually induced to moderate or high levels by adding galactose to the raffinose-containing medium ^98,99^. Transformants were selected based on the TRP1 marker present in the plasmid and were kept in glucose media till the day of the transactivation experiment. Five independent transformants were tested for each p53 mutant protein by preparing fresh patches on selective glucose plates to start liquid cultures for the luciferase assays.

### Luciferase assay

To measure the p53-dependent transactivation of the firefly reporter, we used an optimized low volume liquid culture system ^41,99^. Briefly, independent transformants are resuspended from patches on glucose plates and placed in 200 μl of selective liquid medium containing 2% raffinose as a carbon source within a 96-well plate. 60 μl of cell suspension were then transferred to different 96-well plates and mixed with an equal volume of selective medium containing raffinose as a carbon source, supplemented or not with a desired amount of galactose to induce at different levels the expression of p53. Plates were incubated at 30°C with moderate shaking for 6 hours. In previous experiments, we determined that between 6 and 8 hours in this culture condition, the reporter’s expression of p53 and p53-dependent transactivation reached a peak ^23^. To measure the luciferase activity, 10ul from each well of the cultures were transferred to a white 384-well plate. An equal volume of 2x PLB buffer (Promega) was added to permeabilize the yeast cells. Permeabilization is achieved during a 10 min incubation in a shaker at room temperature. In the meantime, the optical density at 600 nm of each culture was measured in the 96-well plate using a plate reader. Luciferase activity was quantified by adding 10ul of the substrate (Promega) to the permeabilized culture samples in the 384-well plate and measuring luminescence in a plate reader. Luciferase activity is normalized relative to the optical density of the cultures. As a control, transformants with an empty plasmid that do not express p53 were processed in the same manner, and the low-level of p53-independent expression of the firefly reporter was subtracted from the other values.

### Western blotting

To compare p53 protein expression, we developed 3ml liquid cultures in falcon tubes using a selective medium and conditions equivalent to the luciferase assay experiment. 2ml of the cultures were collected in 2ml Eppendorf tubes after six hours and centrifuged at 4000 rpm for less than 2 min to obtain a pellet. Cells were washed once with 5 ml of sterile water and centrifuged again to get a pellet that was then resuspended in 150 μl of 2x PLB buffer. We then added the approximate equivalent of 150 μl of acid-washed, sterile glass beads. Tubes were vortexed at maximum speed for cycles of 30 seconds, followed by one-minute rest on ice. After six such cycles, tubes were spun at maximum speed in a refrigerated microfuge at 4°C for 15 min. The supernatant was then transferred to a new tube, trying to maintain the solution cold. Proteins were quantified by using the bicinchoninic acid assay (Pierce). Western blot was performed with a standard SDS-PAGE protocol in 10% acrylamide gels and loading 10 μg of total protein for each sample. Immunodetection was performed after transferring the protein onto a nitrocellulose membrane and using the DO1 antibody against p53 and an anti PGK1 antibody to detect a reference protein and revealed using a secondary antibody and the ECL detection kit (Amersham). Band intensities were quantified using ImageJ.

