## Supplementary Figures for "Structural Basis of Mutation-Dependent p53 Tetramerization Deficiency"

**SUPPLEMENTARY MATERIAL**

**Supp. Figure 1. Convergence of the MD trajectories.** Values of RMSD as a function of time are plotted for the WT protein and all the mutated forms. Each form rapidly reaches the convergence to structural equilibrium in less than 200 ns.

**Supp. Figure 2. Contact map analysis of WT and mutant p53 TET domain.** Each box shows the average contact map from the different trajectories of the indicated p53 molecules, plotted as a residue index matrix. Mutant p53 forms are expressed as the absolute value of the difference with the WT protein (lower-right corner). The contact maps of R337C, R337D/D352R, R337D, R337$H_{\varepsilon}$, and R337$H_{\delta}$ show significant variations, with the latter presenting the largest divergence. In contrast, D352R and R337$H_{\delta\varepsilon}$ display no substantial deviation from the WT. A threshold of 0.5 was chosen as the cut-off.

**Supp. Figure 3. Transactivation potential of p53 TET mutations at different levels of p53 expression.** Luciferase assays were performed as described in Figure 5 using the indicated reporter strains that are isogenic except for the sequence of the p53 response element that allows p53 to bind to the promoter region upstream of the *Photinus P*yralis reporter and stimulate its transcription. **A**. The p21-5' RE strain is highly responsive to p53. It can register p53-induced transactivation even in cells resuspended directly from glucose plates where the *GAL1* promoter, which controls p53 expression, is at its lowest level of transcription. Using this condition, the difference between p53 alleles is more visible, as the p53 protein levels are limiting. See Figure 5 for a comparison. **B**. Culturing cells in media containing higher levels of galactose (0.032%) stimulates the expression of the luciferase reporter in a p53-dependent manner. However, the difference in activity among the alleles is reduced or no longer appreciable. This is particularly evident with the R337H and D352R in the p21-5' reporter strain and R337H and the double mutant R337D/D352R in the p21-SP2 strain, while D352R continued to exhibit higher activity compared to WT p53. **C.** The reporter strain derived with the spaced p21-5' RE was tested at high galactose (0.128%). The higher relative activity of D352R was confirmed. **D**. To investigate if the effect of D352R was dependent on a generally lower affinity of p53 for its binding sites or more directly to the structural arrangement of RE subunits, we also tested a lower affinity p53 RE derived from the PUMA promoter. In this case, the lower transactivation potential is related to mismatches from the consensus sequence and not to the presence of a spacer between the decameric half-sites. Left and right panels plot data obtained from cultures exposed to 0.008% or 0.032% galactose for six hours, respectively. Results were processed and are plotted as described in Figure 5. * $p<0.05$; ** $p<0.01$; *** $p<0.001$; **** $p<0.0001$, ns = not significant; one-way ANOVA with Dunnett's multiple comparison test.

**Supp. Figure 4. p53 TET mutants are expressed at similar levels in the yeast reporter strains.** Representative western blot (top panel) and quantification of the densitometric scan of the immunodetection comparing the relative expression of WT p53 and the various mutant alleles tested in this study. PGK1 was used as a reference protein. All the p53 tetramerization mutants were expressed at similar levels. We noted a slight decrease in migration in the SDS-PAGE for the double mutant R337D/D352R.

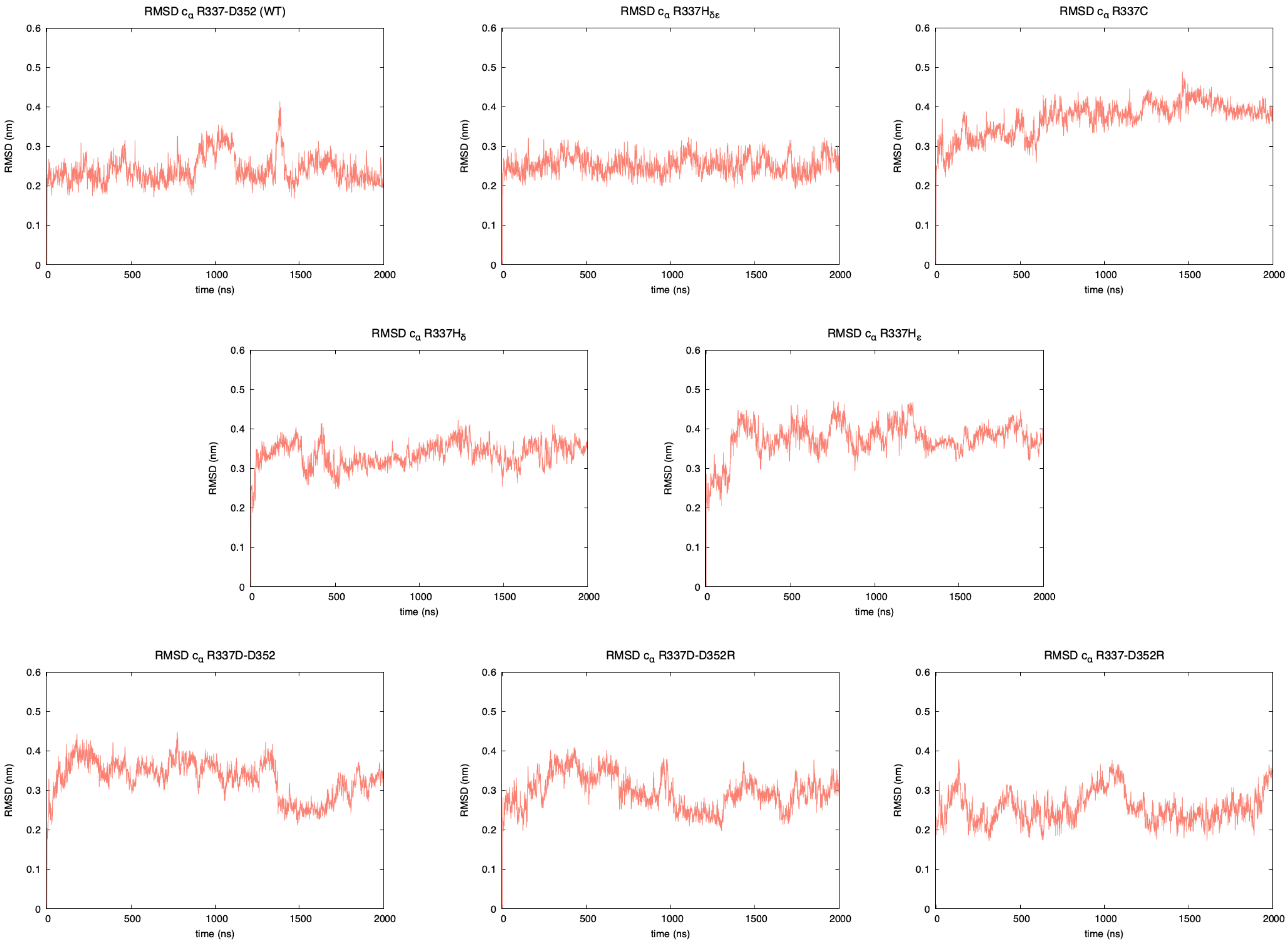


**Supp. Figure 1**

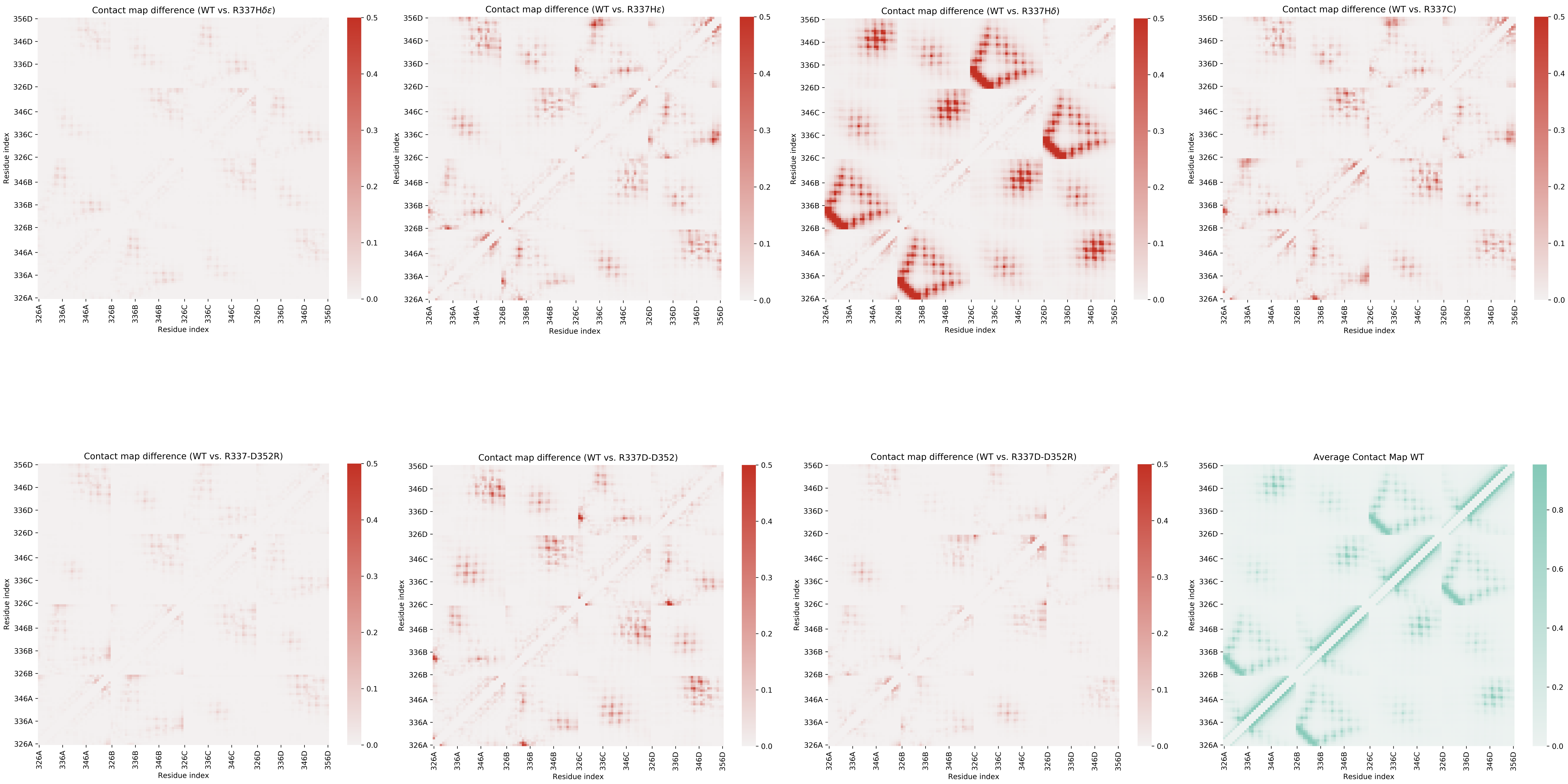


**Supp. Figure 2**

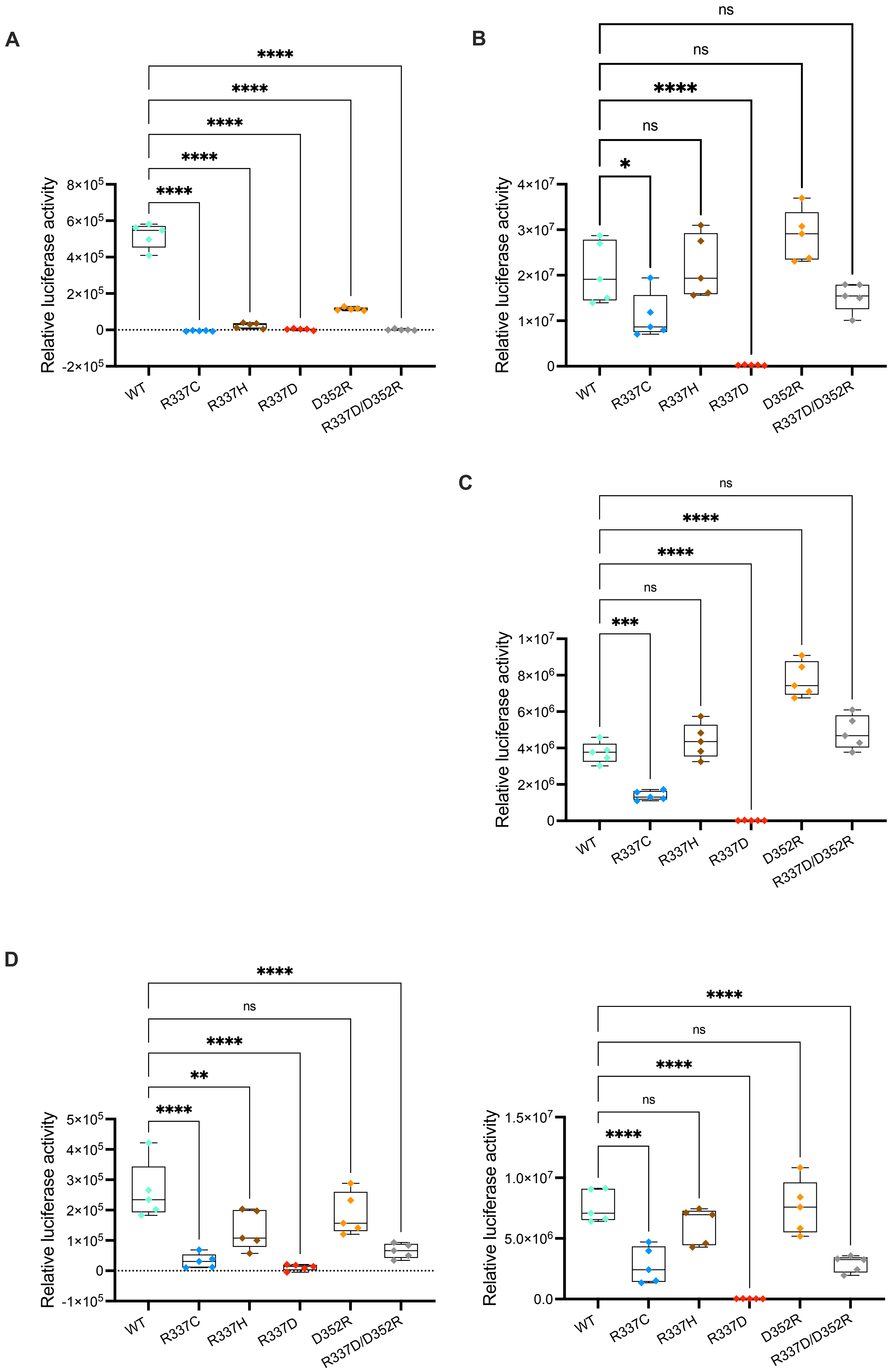


Supp. Figure 3

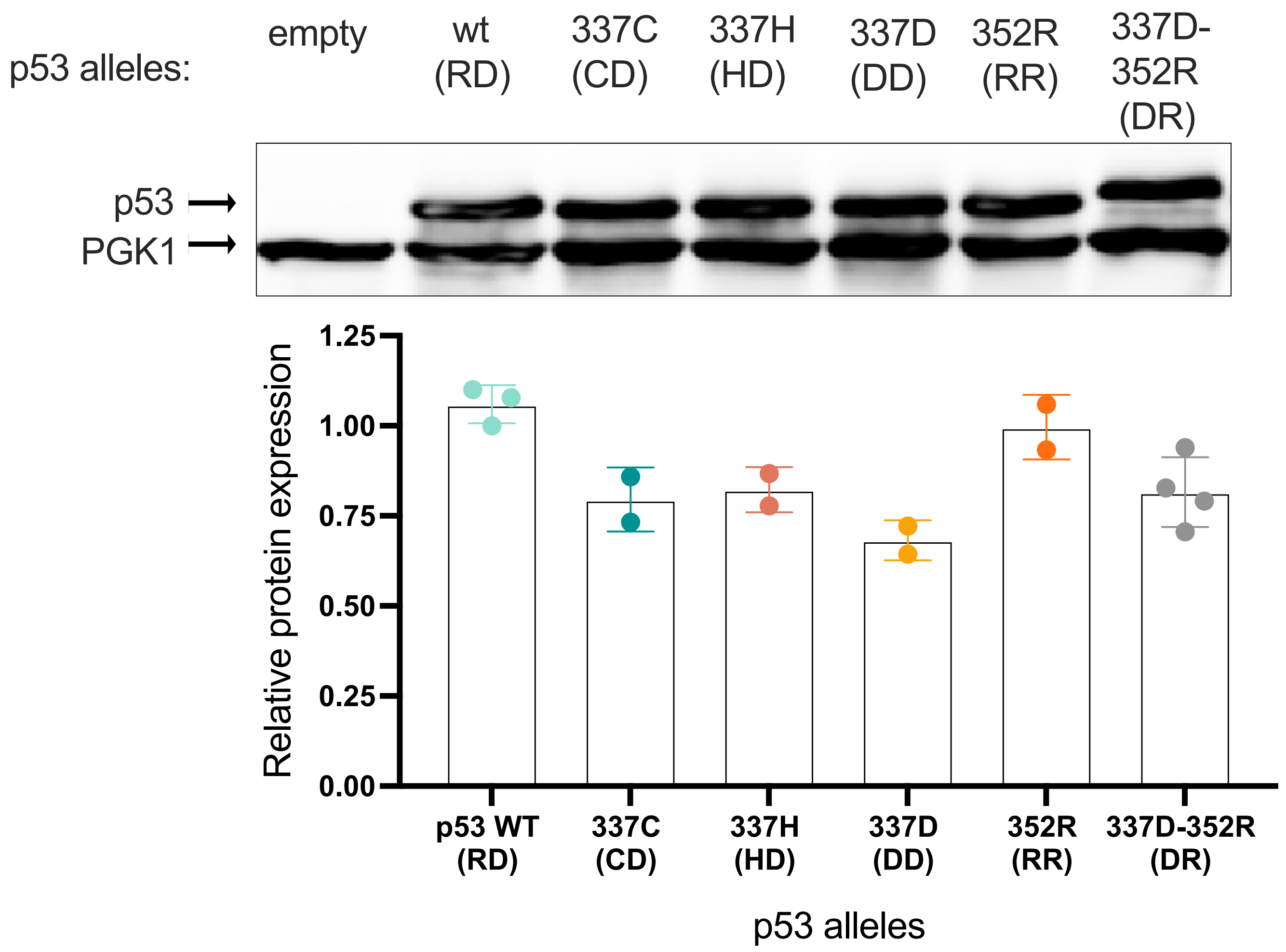


**Supp. Figure 4**
